# Unifying Orientation-Dependent Relaxation and Diffusion Around Axonal Fibers from DTI: Characterizing Fiber-Tract-Specific Anisotropic R_2_ Profiles of the Corpus Callosum

**DOI:** 10.1101/2025.05.02.651936

**Authors:** Yuxi Pang, Rajikha Raja, Wilburn E. Reddick

## Abstract

This work aims to characterize fiber-tract-specific orientation-dependent R_2_ from the seven callosal segments (CC1-CC7) based on human brain Connectome high-resolution DTI datasets. WM voxels from each segment were masked by the thresholds of FA and mode of anisotropy. Encoded in T2W images (b-value = 0), an orientation-dependent R_2_ profile was constructed based on voxel-wise fiber orientations and characterized by a “cone” model. This allowed 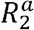, an orientation-dependent or anisotropic R_2_, to be separated from its orientation-independent or isotropic counterpart. Except for 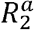, no discernible differences were found for the fits between the two b-values. On average, 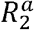 increased from 2.4±0.2 (1/s) to 3.2±0.3 (1/s) as the b-value increased. Furthermore, 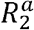 showed an increasing trend from CC1 to CC7 and the open angle α of the cone fluctuated around 65°. As usual, FA was found increasing from CC1 to CC7 and from a higher to a lower b-value. In conclusion, we have shown that fiber-tract-specific Anisotropic R_2_ profiles from the callosal segments can be characterized by a cone model. The proposed method offers a unique opportunity to reevaluate existing clinical DTI studies and optimize new ones for characterizing specific WM tracts in both healthy and diseased subjects.

## 1 INTRODUCTION

In recent years, relaxometry and diffusion have been jointly utilized for microstructure MR imaging of the human brain [1–4]. This combined approach of relaxometry-diffusion offers enhanced imaging specificity for highly ordered microstructures in white matter (WM). This level of enhanced specificity cannot be achieved using either relaxometry or diffusion independently [1]. The biophysical mechanisms underlying these two MR measures are intrinsically linked, yet distinct. More specifically, both relaxometry and diffusion are influenced by the well-known thermally driven Brownian motion of water molecules in biological systems [5,6]. However, the magnetic nuclear relaxation of water protons in biological tissues is predominantly governed by molecular *rotational* diffusion across multiple timescales [7]. In contrast, the conventional diffusion metric derived from diffusion tensor imaging (DTI) is exclusively determined by molecular *translational* diffusion at the cellular level on the timescale of tens of milliseconds [8].

The primary application of microstructure imaging based on relaxometry is the so-called myelin water imaging (MWI) [9,10]. Under normal physiological states in WM, multiple layers of lipid-rich bilayer membranes wrap around an axon [11]. These protective layers, collectively known as myelin sheaths, insulate the axon and thus facilitate rapid electrical impulses along the axon. This rapid electrical conduction can be impaired when myelin sheaths are damaged, leading to compromised sensory, motor, and cognitive functions of the brain [9]. Therefore, MWI can be considered a valuable noninvasive imaging tool for monitoring the integrity of myelin in the human brain. Among the various MWI methods developed so far, the transverse relaxation time *T*_2_ and 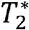 mappings are the most prevalent techniques for compartmentalizing water local environments in WM [12,13,10]. It is worth mentioning that effective transverse relaxation, denoted by 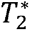, encompasses an irreversible contribution, characterized by *T* _2_.

In biological tissues, water molecules reside in different cellular compartments and interact with a variety of macromolecules, including lipids and proteins [14]. A measured *T*_2_ or 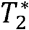 from water molecules may not necessarily correlate with those molecular constituents within a specific cellular water milieu. This is because the observed changes in transverse relaxation times in WM could be further induced by multiple factors, such as iron, myelination, and fiber orientations [15–17]. A recent study on *T*_2_ orientation dependence in the human brain revealed that MWI depends on WM fiber orientations [18]. Specifically, the measured *T*_2_ from myelin water and from intra- and extra-cellular water were both found to be orientation-dependent; moreover, the observed orientation dependence profiles exhibited a similar variation pattern. These interesting findings inarguably cast doubt on the specificity of MWI, as different water compartments should impart unique *T*_2_orientation dependences.

In the past three decades, diffusion MR imaging has been at the forefront of microstructure imaging in the human brain [19]. Because of the spatial resolution gap between standard MR imaging at the millimeter level and the axonal microstructure at the micrometer level, it is inevitable to rely on biophysical modeling to bridge the gap. This modeling allows us to derive microstructural information from macrostructural voxel-wise diffusion measures. The so-called standard model (SM) generalizes various diffusion models that were developed previously for characterizing WM microstructures [8]. It captures the salient features of an elementary axon fiber bundle by two non-exchanging water compartments: an intra-axonal space (IAS) and an extra-axonal space (EAS). The former is characterized by a “stick” diffusion model with an axial diffusivity 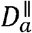 and a signal fraction *f*. In contrast, the latter is described by a “Zeppelin” diffusion model with both axial 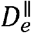 and radial 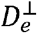 diffusivity, as well as a signal fraction (1- *f*).

According to this SM model, the measured diffusion signal within an image voxel comes directly from a collection of such axon fiber bundles that are orientated differently. This orientation dispersion can be quantified by a fiber orientation distribution (FOD) function. Due to the unfavorable topology of the parameter estimation landscape, it becomes challenging to accurately estimate multiple SM model parameters [20], particularly from noisy diffusion data collected in a clinical setting [8]. As a result, parameter constraints are often introduced to improve the stability of modeling albeit at the cost of reduced model specificity. For instance, 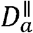 was fixed at 1.7 *µm*^2^/*ms*, and set equal to 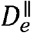 in the widespread neurite orientation dispersion and density imaging (NODDI) [21,22], where the relationship between 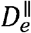 and 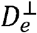 was also constrained.

Even though great progress has been made in uncovering neuronal complex microstructures, a consensus has largely been reached that neither relaxometry nor diffusion can provide an adequate characterization of neuronal microarchitectures in WM [2]. As a result, combined relaxometry-diffusion microstructure imaging has attracted increasing attention in recent years, promising a deeper understanding of neuronal complex microstructures [1]. This insightful understanding stems from the complementary information provided by the joint relaxometry-diffusion approach. Unfortunately, not only does this combined approach extend the time required for imaging data acquisition beyond the limits of standard clinical applications, but it also poses challenges in simultaneously modeling both relaxometry and diffusion phenomena [2].

For instance, compartmentalized anisotropic *T*_2_ in the human brain WM has been recently quantified [23], using different combinations of the *b*-values (ranging from 0 to 6000 s/mm^2^) and the spin echo times (TE) (ranging from 50 to 150 ms). These relaxometry-diffusion data were acquired on a state-of-the-art Connectome 3T MR scanner, equipped with a tiltable RF coil. It was found that the water signal from EAS manifested a stronger *T*_2_ orientation dependence effect than that from IAS. Moreover, the measured *T*_2_ values from EAS were significantly smaller than that from IAS, which is in accordance with the literature [24]. However, these interesting findings appear to conflict with the SM model, where IAS is considered more anisotropic than EAS [8]. Furthermore, the reported orientation-dependent *T*_2_ profiles from IAS are inconsistent with those found in neonatal WM [23,25,26], where anisotropic *T*_2_ contributions from EAS should be negligible due to incomplete myelination. It is worth mentioning that myelin water, which plays a critical role in MWI [18,27], has not been included in the SM model due to its short *T*_2_ relative to the typical TE used in clinical applications [8].

Given the nature of multidimensional data in combined relaxometry-diffusion imaging, it has become increasingly critical to identify the most relevant degrees of freedom from any developed practical biophysical models [19]. This identification is essential for mitigating the abovementioned challenges associated with not only extensive data acquisitions but also sophisticated biophysical models. In this context, anisotropic relaxometry and anisotropic diffusion should be considered directly interconnected for a typical axonal fiber bundle in WM [17,28]. Consequently, WM microstructures could be better characterized by the most relevant metrics derived from anisotropic relaxometry and anisotropic diffusion. To achieve this goal, it becomes imperative to fully disentangle the anisotropic components from their isotropic counterparts in relaxometry and diffusion.

Recently, a generalized magic angle effect function has been proposed for characterizing anisotropic transverse *R*_*2*_ relaxation [28,17,29], which is orientation-dependent in highly organized biological tissues. This specific function was derived from a so-called cone model, which describes a specific distribution of the residual dipolar interaction vectors stemming from rotationally restricted water or other molecules. The derived model parameters could be used to quantify the extent of myelination due to normal brain development or aging, and pathological changes such as demyelination in WM diseases [30,25,31]. Potentially, these derived relaxation metrics could also be useful for characterizing the axonal loss concurrent with pathological changes. With respect to those metrics derived from translational diffusion measurements, anisotropic *R*_*2*_ boasts more specificity as it is directly linked to the underlying WM microstructures [28,17].

With these insights in mind, the aim of this work is to introduce an efficient method for extracting and characterizing fiber-tract-specific anisotropic *R*_*2*_ from DTI. The proposed method was applied to the segmented tracts passing through the corpus callosum (CC) in the human brain, shedding more light on the underlying WM microstructural alterations across various callosal segments.

## 2 THEORY

### 2.1 A diffusion-weighted spin-echo signal formula

The signal intensity of an image voxel, denoted as *S*_*b* ≠0_, from diffusion-weighted (DW) spin-echo (SE) image in an isotropic medium can be adequately described by Eq. 1 [5]:

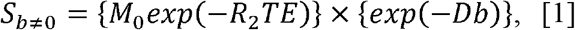

where *M*_0_, *R*_2_, *TE*,*D* and *b* denote, respectively, a constant affected by both intrinsic (e.g., local spin density) and extrinsic (e.g., specific hardware settings) tissue factors, a transverse relaxation rate, a spin echo time, an effective translational diffusion coefficient, and a diffusion weighting factor. The diffusion weighting factor, *b*, depends on diffusion gradient strength, duration, and separation between a pair of such gradients. This signal formula in Eq. 1 encapsulates the two essential aspects of molecular Brownian motions, i.e., the rotational and the translational, which are respectively included in the first and second curly brackets.

In highly organized biological tissues, the transverse relaxation is predominantly governed by intramolecular dipolar interactions of water protons [29,28]. These dipolar interactions are modulated by molecular rotational diffusion on various timescales ranging from “fast” pico-nanoseconds to “slow” micro-milliseconds [7,32]. When the diffusion gradient is absent, the signal intensity, denoted as *S*_*b* = 0_, restores to that from the standard SE image in highly organized biological tissues, as expressed by Eq. 2:

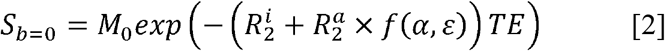

Herein, *R*_*2*_ has been separated into an isotropic 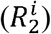 and an anisotropic 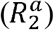 component for white matter (WM) [17,28]. The orientation ( *ε*) dependence of the latter is characterized by the function *f(α*,*ε)* as written by Eq. 3:

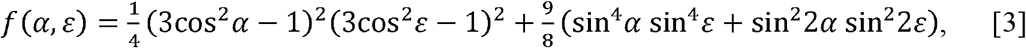

It should be emphasized that the function *f(α*,*ε)*was derived from an axially symmetric or “cone” model as schematically depicted in Fig. 1a for residual dipolar interaction vectors distributed around an axon fiber [17,28]. Herein, *α* and *ε)*denote, respectively, an open angle of the cone and an angle of the axon fiber relative to the direction of a static magnetic field *B*_0_.

**Fig. 1.**
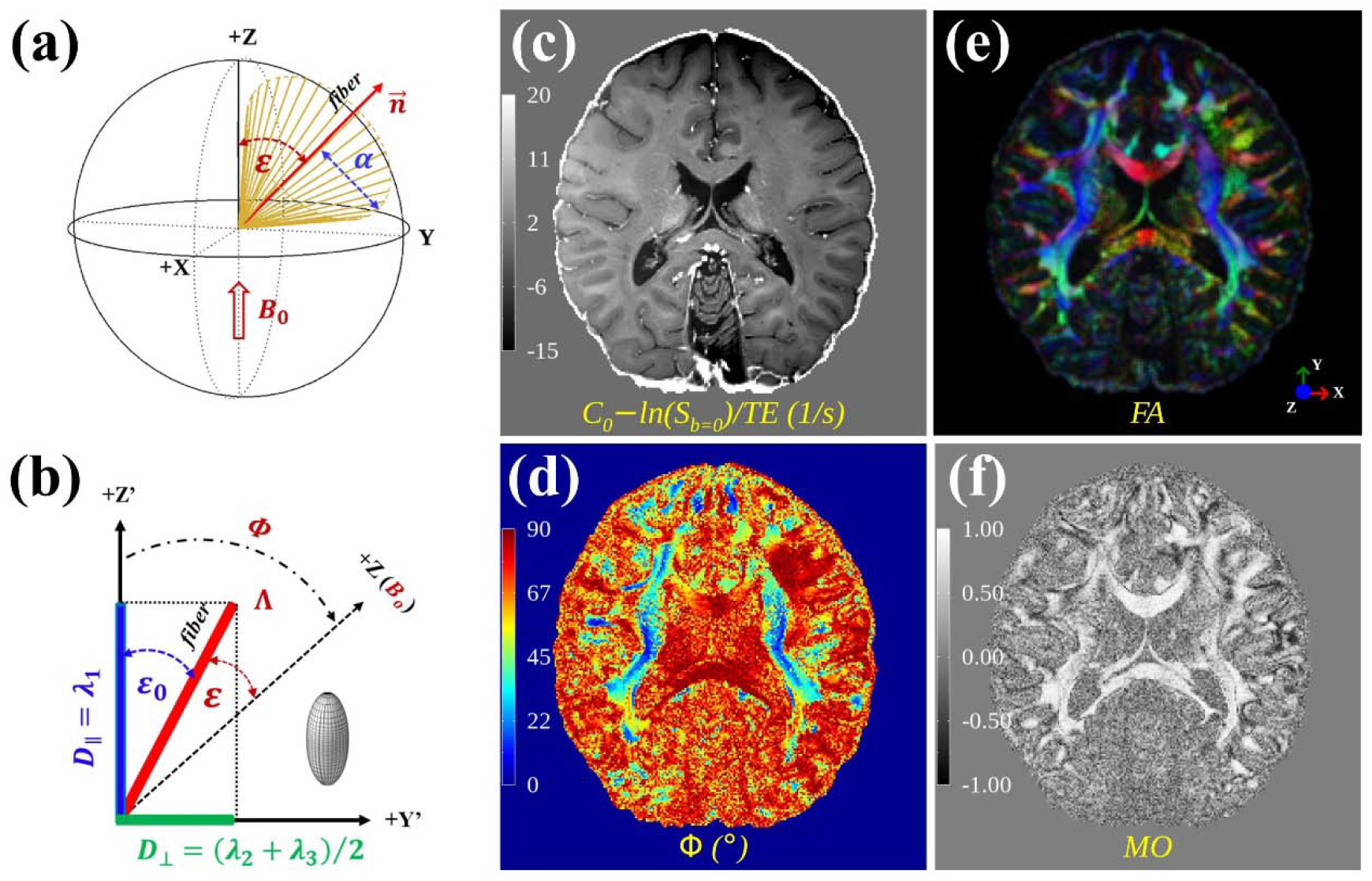
Schematics of an orientation-dependent R_2_ “cone” model (a) and the direction of a “cigar-like” diffusion tensor relative to B_0_ (b). The figure also includes four parametric maps derived from a high-resolution DTI dataset, namely, (c)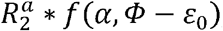, (d) principal generated with C_0_ arbitrarily set to 96 (1/s) for demonstration purposes. diffusivity direction, (e) fractional anisotropy, and (f) the mode of anisotropy. Note: Fig. 1c was

If both 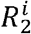 and *M*_*0*_ are assumed constant in WM, both 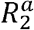 and *α* can be determined given the knowledge of *ε*, according to Eqs. 2 and 3. The assumptions involved have been supported by previously published experimental results [17]. Specifically, in WM across the whole brain, the observed variations of *R*_*1*_ or *M*_*0*_ with fiber orientations at 3T were significantly smaller compared to those of transverse relaxation rates [33,34]. Furthermore, the fitted model parameters for the orientation-dependent *R*_*2*_ profiles derived from both T2-weighted (T2W) signals and *T*_2_ values are essentially indistinguishable, suggesting that the aforementioned assumptions are valid [17]. These assumptions will become even more robust for a specific segmented WM tract due to its local nature. Herein, the variation of *R*_*1*_ could be considered as a proxy of the variation of 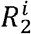, as both relaxation metrics are mostly governed by water molecular reorientations on the timescales of nanoseconds or less [7,32,35].

### 2.2 A voxel-based anisotropic diffusion direction

The translational diffusion coefficient, *D*, of water molecules in an isotropic medium can be calculated based on Eq. 1, i.e., *D= (* 1/ *b)* ln (*S*_*b* = 0;_/ *S*_*b* ≠0_) in other words, only a single non-zero value is required to derive such a scalar diffusion metric. In highly organized tissues such as WM in the brain, however, the translational diffusion of water molecules becomes anisotropic or dependent on orientation. In this scenario, *D* can be characterized by a second-rank symmetric tensor that has six independent elements [36]. Consequently, this symmetric diffusion tensor can be determined using multiple DW images in which the same diffusion gradient (*b* ≠0) is repeatedly applied along at least six non-collinear directions. This experimental framework underlies the fundamental physical basis for DTI.

Once the diffusion tensor is determined in the laboratory reference frame (LRF), where the direction of *B*_0_ coincides with +Z axis, it can be mathematically transformed into a diagonalized matrix in the principal axis system (PAS). This unique system is characterized by three eigenvalues in descending order, i.e., *λ*_1_ > *λ*_2_ > *λ*_3_ The corresponding three eigenvectors (*ê*_*1*_, *ê*_*2*_, *ê*_*3*_) are also determined, representing the directions of the primary (*λ*_1_), secondary (*λ*_2_), and tertiary (*λ*_3_) diffusivities in LRF, respectively. As a result, a series of rotationally invariant quantities can be defined based on these three eigenvalues. For instance, the mean diffusivity (MD) and fractional anisotropy (FA) are calculated by ∑ *λ*_*i*_/3 with *i*= 1,2,3 and by 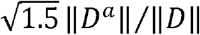, respectively. Note, the magnitudes of an anisotropic and an effective diffusion tensor are respectively given by 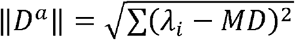 and by 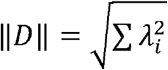.

In the literature, the direction of *D*^*a*^ or *D* was not explicitly defined; nonetheless, both are theoretically demonstrated to be collinear [36]. Intuitively, this direction should align with the direction (*ε*) of an effective axon fiber. However, in the past, the direction (Φ) of *λ*_1_ has been *assumed* to be a surrogate for *ε*. This practice seems to be inconsistent with the definition of FA, as all three eigenvalues are included in determining the magnitude of an anisotropic diffusion tensor *D*^*a*^. Follow the same logic as the FA definition, the direction of *D*^*a*^ should also be determined by all three eigenvalues unless *λ*_1_ ≫ *λ*_2_ > *λ*_3_— a condition that is most unlikely in WM. In our recent work [17], *ε* has been demonstrated to deviate from Φ by an offset angle *ε*_0_, i.e., *ε* = Φ − *ε*_0_, for an axially symmetric prolate (i.e., *λ*_1_ >*λ*_2_ ≈*λ*_3_) diffusion tensor, as schematically depicted in Fig. 1b. Φ

### 2.3 A tract-specific orientation-dependent *R*_2_ profile

In a logarithmic scale, Eq. 2 can be recast into Eq. 4:

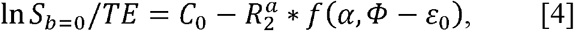

where *C*_0_ denotes an unknown constant represented by 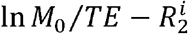. If a specific WM tract can be segmented from the whole brain based on DTI, the relevant voxels from this tract will be pooled and binned as previously demonstrated in the literature [23,18]. Subsequently, a tract-specific orientation-dependent *R*_2_ profile (i.e., *C*_0_ – ln *Sb*=0/*TE*,) can be established, and the four model parameters (i.e., 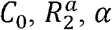, and *ε*_0_) can be extracted based on Eq. 4.

## 3 METHODS

### 3.1 A high-resolution DTI dataset in public domain

To corroborate the proposed theoretical framework, this work utilized publicly available Connectome DTI datasets of an in vivo human brain at 760 *μm*^3^ isotropic resolution [37]. These datasets, acquired with two non-zero *b*-values (1000 and 2500 s/mm^2^) and TE = 75 ms, had been preprocessed to correct for image artifacts arising from signal drifts, susceptibilities, eddy currents, gradient nonlinearity induced distortions, and participant involuntary head motions. Six (*b*=1000 s/mm^2^) and twelve (*b*=2500 s/mm^2^) preprocessed data subsets were analyzed separately using FSL DTIFIT software [38].

The output parameters from DTIFIT included three eigenvalues *(λ*_*1*_, *λ*_2,_ *λ*_3_), three eigenvectors *(ê*_*1*_, *ê*_2,_ *ê*_3_), FA (fractional anisotropy), MO (mode of anisotropy) [39], and T2W signal s_*b*=0_. The direction (*Φ*) of principal diffusivity *λ*_*1*_ was determined by the primary eigenvector *ê*_*1*_ through the relationship [18]: cos *Φ =ê*_1_.*B*_0_/‖ *ê*_*1*_.*B*_0_‖. Four parametric maps are illustrated in Fig. 1: *C*_*0*_ − ln (*S*_*b*=0_)/*TE*, (Fig. 1c), *Φ* (Fig. 1d), colored coded FA (Fig. 1e), and MO (Fig. 1f). These maps were derived from the data subsets with *b*=1000 s/mm^2^, with *C*_0_ arbitrarily set to 96 (1/s) for demonstration purposes. This assumed *C*_0_ value approximates those fitted from the orientation-dependent *R*_2_ profiles of various callosal segmentations (see Table 1).

**Table 1.** Derived relaxometry metrics for seven callosal segments (CC1-CC7) from diffusion datasets with b-values of 1000 and 2500 (s/mm^2^). The fitted constant C_0_ (1/s), anisotropic 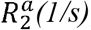, open angle α(°)of the “cone” mode, phase shift ε_0_ (°), and goodness-of-fit R^2^ (%) are listed from the 3^rd^ to the 7^th^ column, respectively. All metrics are given by mean±std, and the average (CC AVE) across the entire corpus callosum is also provided.

| $b$ | Name | $C_0$ (1/s) | $R_2^a$ (1/s) | $\alpha$ ($^\circ$) | $\varepsilon_0$ ($^\circ$) | $R^2$ (%) |
| --- | --- | --- | --- | --- | --- | --- |
| 1000 ($\text{s/mm}^2$) | CC1 | 98.0±0.2 | 1.2±0.3 | 59.1±6.7 | 25.5±1.6 | 84.1 |
| | CC2 | 98.5±0.2 | 1.0±0.2 | 62.4±2.3 | 1.4±3.2 | 84.8 |
| | CC3 | 97.3±0.4 | 1.0±0.7 | 74.8±2.8 | 15.6±13.2 | 11.7 |
| | CC4 | 98.3±0.4 | 3.3±0.7 | 63.1±3.1 | 26.8±1.2 | 91.4 |
| | CC5 | 98.7±0.1 | 2.4±0.3 | 54.3±7.8 | 27.7±1.2 | 87.8 |
| | CC6 | 102.0±0.1 | 6.3±0.2 | 69.0±0.3 | 10.0±1.4 | 99.3 |
| | CC7 | 101.3±0.2 | 3.6±0.3 | 61.3±1.6 | 10.3±1.2 | 96.6 |
| | CC AVE | 98.9±0.1 | 2.4±0.2 | 64.6±1.1 | 19.8±1.0 | 97.7 |
| 2500 ($\text{s/mm}^2$) | CC1 | 97.2±0.3 | 1.2±0.1 | 50.9±9.2 | 34.2±0.9 | 90.6 |
| | CC2 | 98.9±0.1 | 2.0±0.1 | 65.4±0.6 | 0.0±0.0 | 93.6 |
| | CC3 | 97.3±0.2 | 0.8±0.4 | 74.6±4.2 | 2.7±14.2 | 28.6 |
| | CC4 | 99.9±0.5 | 5.9±0.8 | 68.6±1.0 | 24.0±1.8 | 90.8 |
| | CC5 | 98.9±0.3 | 3.0±0.7 | 58.9±5.6 | 27.3±1.1 | 90.1 |
| | CC6 | 101.9±0.2 | 6.9±0.3 | 68.3±0.4 | 13.7±1.2 | 98.7 |
| | CC7 | 101.6±0.3 | 4.5±0.3 | 63.4±1.0 | 7.7±1.3 | 97.2 |
| | CC AVE | 99.1±0.2 | 3.2±0.3 | 66.3±0.8 | 19.4±1.1 | 97.7 |

### 3.2 Callosal segmentation

To enhance the imaging specificity, we focused on the segments of corpus callosum — one of the major WM tracts in the human brain. A fast and accurate WM tract segmentation tool, TractSeg [40], was employed to partition the CC into seven subregions [41]: rostrum (CC1, red), genu (CC2, blue), rostral body (CC3, green), anterior mid body (CC4, pink), posterior mid body (CC5, yellow), isthmus (CC6, cyan), and splenium (CC7, purple), as depicted in Fig. 2.

**Fig. 2.**
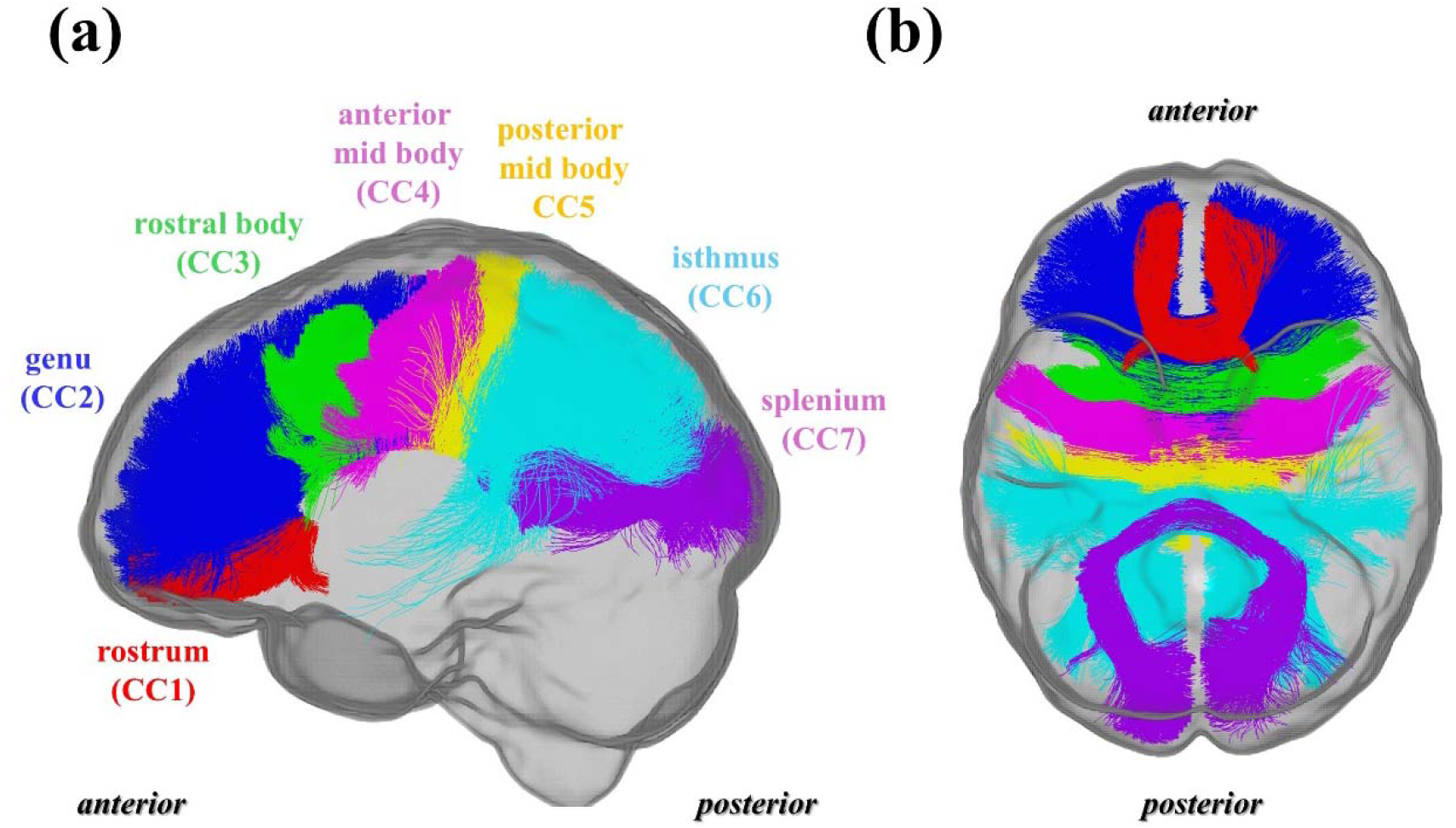
Sagittal (a) and axial (b) views of the fiber tracts pass-through corpus callosum, showing seven segmented tracts: Rostrum (CC1, red), Genu (CC2, blue), Rostral body (CC3, green), Anterior mid body (CC4, pink), Posterior mid body (CC5, yellow), Isthmus (CC6, cyan), and Splenium (CC7, purple).

More specifically, the preprocessed DTI datasets were initially aligned with the Montreal Neurological Institute (MNI) space using the standard FA template from FSL. These aligned datasets were used as input for the TractSeg tool to generate the FOD peak images, thereby pinpointing the WM tracts of interest. These selected tracts in the MNI space, which served as WM voxel masks, were then transformed back to their original space.

The masked voxels from these seven tracts were further culled by the thresholds of 0.5 < FA < 0.9 and 0.5 < MO < 1.0 [17]. Consequently, each isolated voxel from the CC could be well characterized by an approximately prolate, cigar-shaped diffusion tensor. The measured T2W signal *s*_*b*=0_ (in a logarithmic scale) from these selected voxels in the seven segmented tracts were separately sorted, based on their respective *Φ*, and then averaged into 30 different bins spanning from 0° to 90°.

### 3.3 Nonlinear least-squares curve-fitting

To characterize an orientation-dependent *R*_2_ profile, the sorted and binned *S*_*b*=0_ values were fitted to the model characterized by Eq. 4. The unweighted nonlinear least-squares fits were performed using an Interactive Data Language (IDL) programming script from the public domain (http://purl.com/net/mpfit). There were four parameters in this model, which were constrained to the following ranges: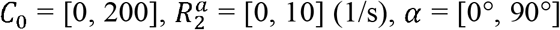, and *ε* _0_= [0°, 45°]. The uncertainties for the fits were adjusted so that the reduced *χ* ^2^ became unity, and the goodness-of-fit was represented by an adjusted *R*_2_ in terms of percent (%). Unless stated otherwise, the standard deviations of the presented data were indicated by either error bars or the half-widths of shaded ribbons. All data visualization and analysis were conducted using in-house software developed in IDL 8.9 (Harris Geospatial Solutions, Inc., Broomfield, CO, USA).

## 4. RESULTS

### 4.1 Orientation-resolved callosal voxel counts

Fig. 3 presents the numbers of orientation-resolved voxels within individual segmented tracts from the corpus callosum (CC) for data subsets with *b*-values of 1000 s/mm^2^ (Fig. 3a) and 2500 s/mm^2^ (Fig. 3b). In general, the voxel number decreased with either a lower orientation angle or a higher *b*-value, consistent with the literature [42,17]. Among the individual tracts, the most populated were the genu (CC2) and the isthmus (CC6), while the rostrum (CC1) was the least.

**Fig. 3.**
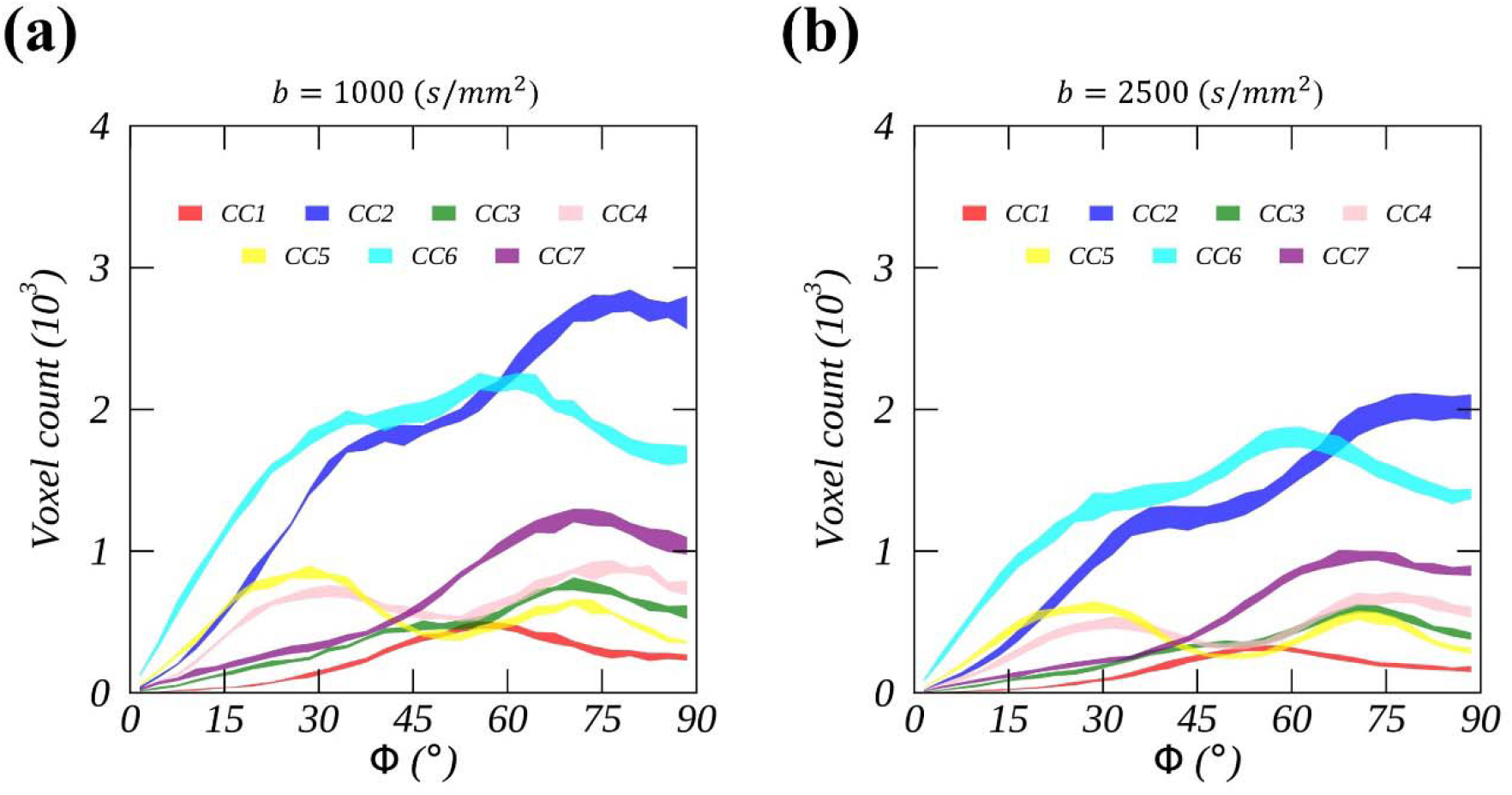
The orientation-resolved voxel numbers for seven callosal fiber tracts derived from diffusion datasets with b-value (s/mm^2^) of 1000 (a) and 2500 (b). These fiber tracts are as follows: Rostrum (CC1, red), Genu (CC2, blue), Rostral body (CC3, green), Anterior mid body (CC4, pink), Posterior mid body (CC5, yellow), Isthmus (CC6, cyan), and Splenium (CC7, purple). Note, the width of the ribbon indicates the uncertainty of the voxel counts.

### 4.2 Measured and fitted orientation-dependent *R*_2_ profiles

Fig. 4 illustrates the measured (represented by shaded ribbons) and fitted (depicted by red lines) orientation-dependent *R*_2_ profiles for segmented tracts from CC1 to CC7 (Fig. 4a-g), derived from data subsets with *b*=1000 s/mm^2^. An average from the entire CC is also included (Fig. 4h). In the same arrangements, the measures and fits from data subsets with *b*=2500 s/mm^2^ are shown in Fig. 5. Table 1 lists the fitted model parameters, along with the goodness-of-fit *R*_2_. In a broad sense, the proposed model (i.e., Eq. 4) can reasonably explain the measured data, as quantified by *R*_2_> 84%. However, this was not the case for the rostral body (CC3) tract (*R*_2_< 30%). Currently, we do not know how to correctly interpret the observed CC3 profiles. Consequently, the measurements and fits from CC3 are not further discussed in the subsequent analysis unless otherwise specified.

**Fig. 4.**
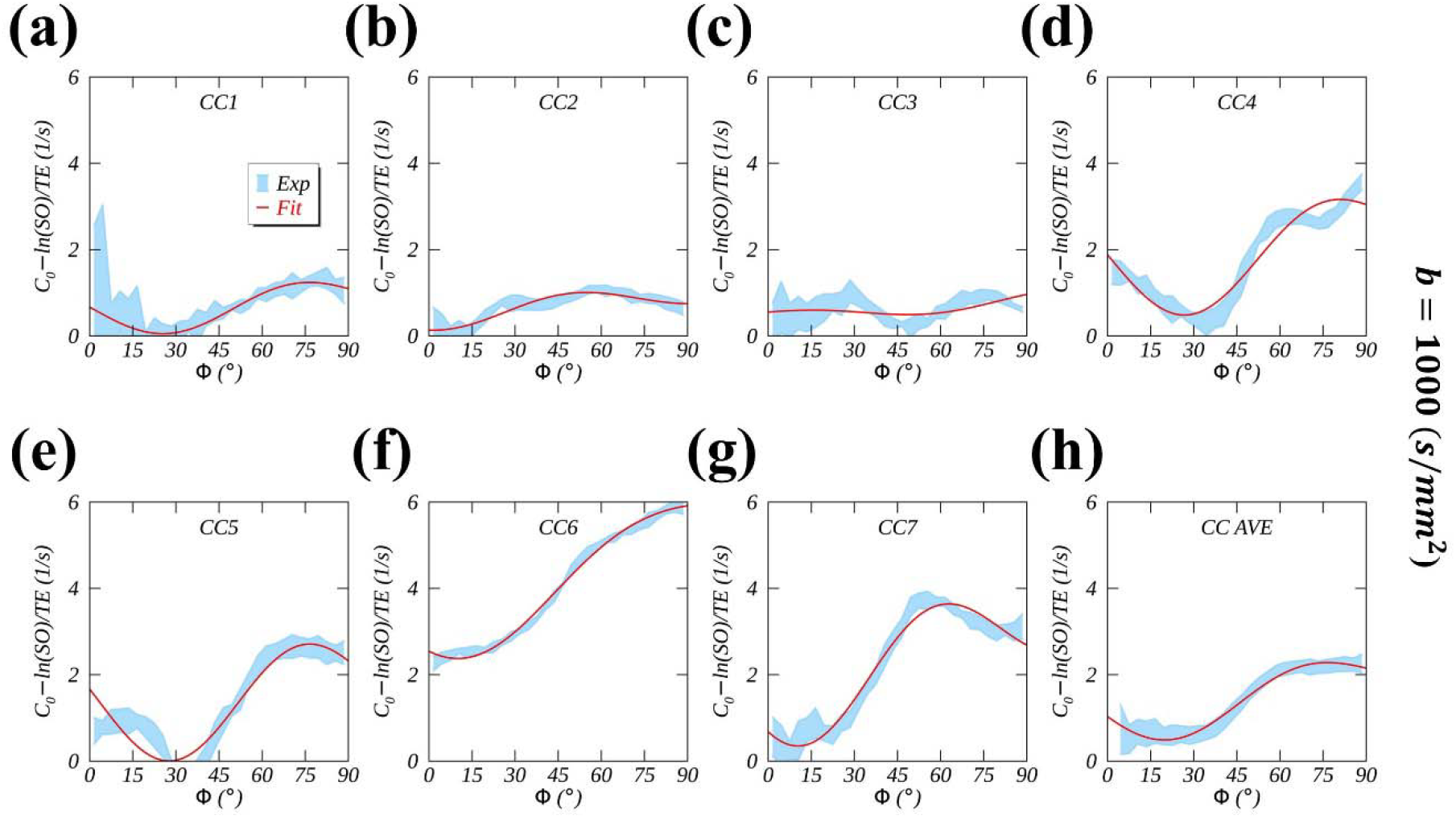
Measured (denoted by ribbons) and fitted (denoted by red lines) orientation (Φ)- dependent R_2_ profiles for segmented callosal fiber tracts from CC1 (a) to CC7 (g). These the average orientation-dependent R_2_ profiles across the entire corpus callosum (h). profiles were derived from diffusion datasets with a b-value of 1000 (s/mm^2^). Also included are

**Fig. 5.**
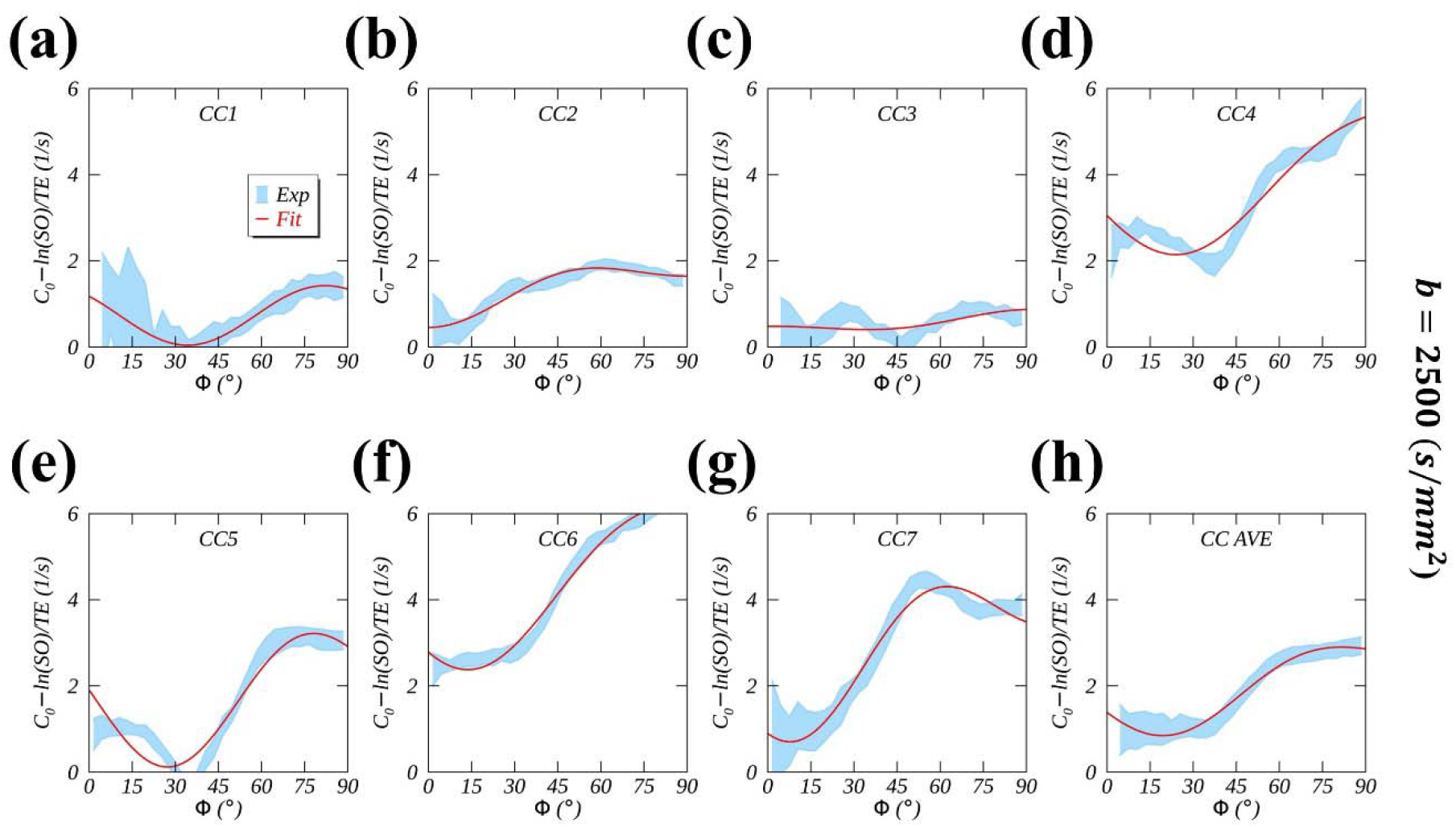
Measured (denoted by ribbons) and fitted (denoted by red lines) orientation (Φ)- dependent R_2_ profiles for segmented callosal fiber tracts from CC1 (a) to CC7 (g). These the average orientation-dependent R_2_ profiles across the entire corpus callosum (h). profiles were derived from diffusion datasets with a b-value of 2500 (s/mm^2^). Also included are

### 4.3 Callosal diffusion and relaxation metrics

Fig. 6 compares the derived diffusion (Fig. 6a-b) and relaxation (Fig. 6c-f) metrics across the callosal segments for *b*-values of 1000 s/mm^2^ (red circles) and 2500 s/mm^2^ (blue squares). For the derived diffusion metrics, FA showed an increasing trend from the anterior CC1 to posterior CC7 and it increased significantly when moving from a higher to a lower *b*-value (Fig. 6a). For instance, a pair of FA values were, respectively, 0.566±0.018 (CC1) vs. 0.594±0.008 (CC7) for *b*=1000 s/mm^2^, and 0.548±0.009 (*b*=2500 s/mm^2^) vs. 0.596±0.006 (*b*=1000 s/mm^2^) for CC5. These results are in good agreement with the literature [43–45]. As shown in Fig. 6b, MD decreased with a higher *b*-value and exhibited a slightly increasing trend from CC4 to CC7, particularly with a lower *b*-value. Interestingly, a recent study found opposite and same trends for MD and FA, respectively, in the comparable callosal segments at 7 T [46].

**Fig. 6.**
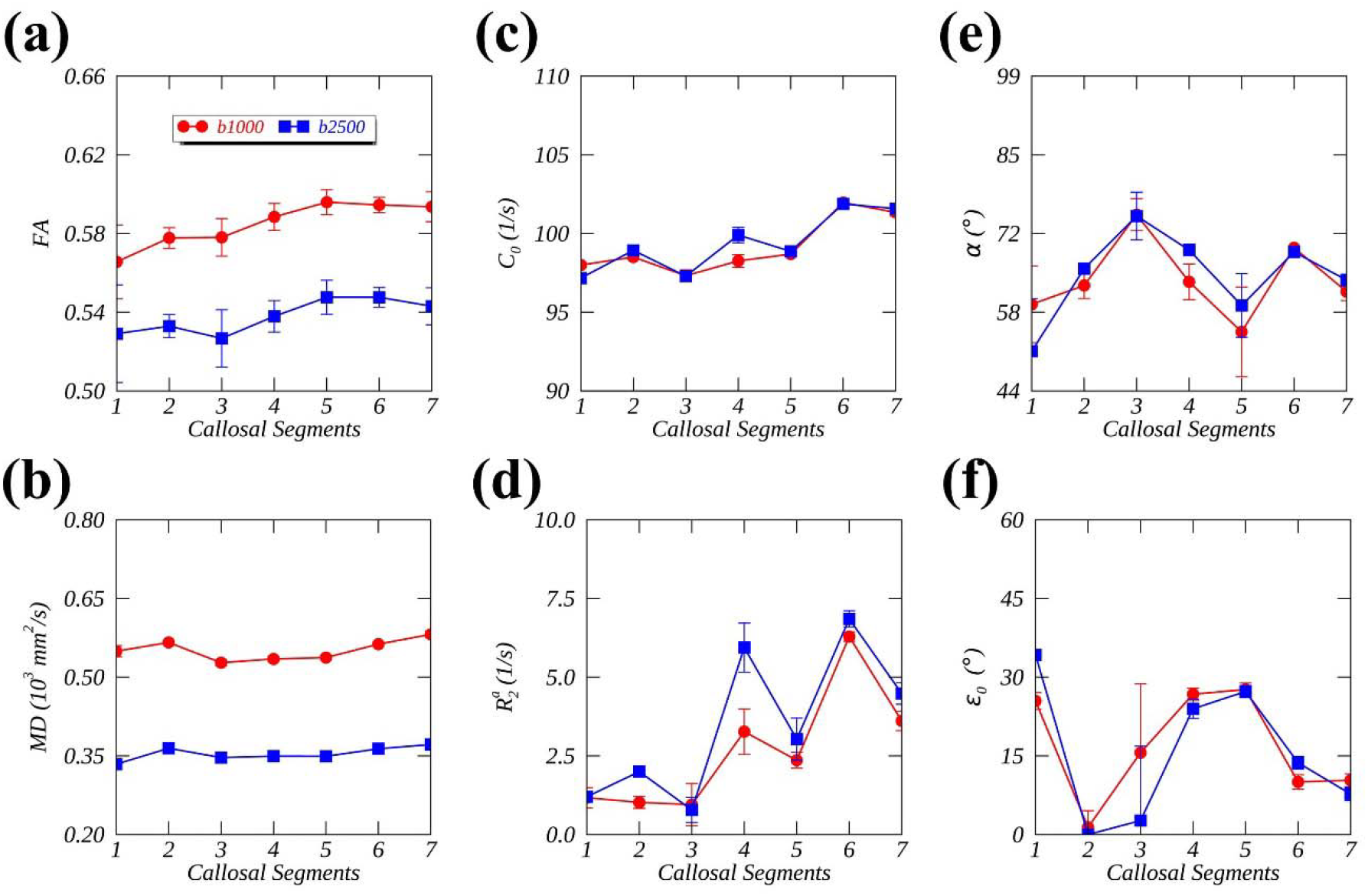
Diffusion (a-b) and relaxometry measures (c-f) for seven callosal segments derived from diffusion datasets with a b-value of 1000 (red symbols) or 2500 (s/mm^2^) (blue symbols). of Specifically, diffusion metrics include (a) fractional anisotropy and (b) mean diffusivity (mm^2^/s), and relaxometry metrics are (c) constant C_0_ (1/s), (d) anisotropic 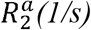, (e) open angle α(°) the “cone” mode, and (f) phase shiftε_0_ (°).

On the other hand, for the derived relaxation metrics between two *b*-values, no discernible differences were found except for 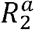 (Fig. 6d). On average, as listed in Table 1, 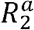 increased from 2.4±0.2 (1/s) to 3.2±0.3 (1/s) as the *b*-value increased, which aligns with the previous findings from the whole brain [17]. From the callosal segments CC1 to CC7, both *C*_0_ (Fig. 6c) and 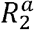 (Fig. 6d) showed an increasing trend. The derived *α* (Fig. 6e) fluctuated around 65°; however, the derived *ε*_0_ (Fig. 6f) showed an apparently decreasing trend when CC2 and CC3 were excluded. It should be emphasized that the relative changes in *C*_0_, which ranged from about 97 (1/s) to 102 (1/s), were approximately 5% and thus considered negligible when compared to those in 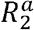, which ranged from about 1.0 (1/s) to 7.0 (1/s), representing a 600% increase. These findings suggest that the previous assumptions about the constant 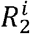 and *M*_0_ were justified in the human brain WM.

## 5 DISCUSSION

This work introduces an efficient method for characterizing fiber-tract-specific anisotropic *R*_2_ from DTI. The inherent anisotropic relaxometry information, encoded in images with a *b*-value of zero, can be successfully recovered. This information is specific to the axonal fibers in the human brain. Our study significantly simplifies previous joint relaxometry-diffusion schemes, which are not only time-consuming but also complex. This achievement stems from focusing on the most relevant information in an anisotropic environment within WM.

### 5.1 Identifying the relevant degrees of freedom in relaxometry and diffusion modeling

Magnetic nuclear relaxation comprises longitudinal and transverse components [6]. In biological systems, these two relaxation processes are predominantly governed by intramolecular dipolar interactions on multiple timescales [7]. While both longitudinal and transverse relaxation rates, *R*_1_ and *R*_2_, are determined by stochastic modulations of dipolar interactions on the “fast” timescales spanning from picoseconds to nanoseconds, *R*_2_ is further enhanced by some molecular interactions on the “slow” timescales of microseconds to milliseconds. Examples of these slow interactions include the magic angle effect [32] and chemical exchange effect that includes diffusion through varying microstructural susceptibilities in biological tissues [47,48].

These slow molecular interactions are often associated with some specific molecular constituents in biological tissues, for instance, collagen [32] and glycosaminoglycan [48] in articular cartilage. Consequently, separating molecular interactions on the slow timescales from those on the fast timescales, for instance, through *R*_1*ρ*_ dispersion or orientation-dependence, will provide exclusive imaging specificity. This work has exploited the phenomenon of *R*_2_ orientation dependence in WM, successfully separating an anisotropic 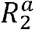 from its counterpart 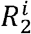, thus specifically quantifying the extent of (de)myelination [17,28].

On the other hand, the amplitude and direction of anisotropic diffusion from DTI were not consistently defined from the beginning [36]. The normalized amplitude, i.e., fractional anisotropy, was calculated from all three eigenvalues; in contrast, the direction was normally *assumed* to be the direction (Φ) of principal diffusivity *λ*_1_. This assumption becomes reasonable only for some specific diffusion tensors when *λ*_1_ ≫ *λ*_2_ >*λ*_3_. Recently, we have demonstrated that the direction of anisotropic diffusion should also be determined by all three eigenvalues [17]; otherwise, the measured orientation-dependent *R*_2_ profiles, guided by Φ, would manifest a phase shift *ε*_*0*_.

In the proposed method, the most relevant diffusion information comes from the direction ( *ε*) of anisotropic diffusion (see Fig. 1b) along an axon fiber, around which rotationally restricted water molecules are arranged. Although DTI contains not only translational diffusion but also rotational diffusion information encoded in T2W images with a *b*-value of zero, the latter has never been exploited in the past except as a normalization reference. The present work has successfully recovered the specific information from orientation dependent *R*_2_. It is worth mentioning that the diffusion metrics derived from DTI heavily depend on the choices of *b*- values, and the potential kurtosis effect [49] could confound the reported findings with *b*=2500 s/mm^2^. Nonetheless, this work has clearly demonstrated that an insightful theoretical framework can significantly simplify practical applications [19].

### 5.2 Masking white matter voxels from the whole brain

The reliability of an orientation-dependent *R*_2_ profile largely depends on accurately separating WM voxels from the whole brain tissue. In the past, a predefined FA threshold was commonly utilized to identify WM image voxels from a co-registered 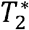 parametric map [42]. Alternatively, a WM mask can also be generated from high-resolution *T*_1_-weighted images [18]. This latter approach can minimize potential registration bias, especially when segmentation is performed using advanced tools like FreeSurfer [50], which are typically optimized for *T*_1_-weighted contrast. It is possible that the separated voxels may not originate from WM partially due to misregistration between diffusion weighted and 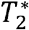 or *T*_1_-weighted image. Furthermore, a specific FA value may represent either a prolate or an oblate shape of the diffusion tensor. In the present work, both *T*_2_ and diffusion weighted images originate from the same DTI acquisition, thus eliminating potential errors induced by misregistration. Moreover, an additional constraint, i.e., the mode of anisotropy, was used to ensure the uniqueness of the diffusion tensor shape determined by FA. Even with these stringent selection criteria, the segmented CC3 exhibited an unusual orientation-dependent *R*_2_ profile that could not be characterized by the proposed model. However, this specific failure should not lead to questioning the credibility of the proposed model, as it has been thoroughly validated in literature and worked well for the whole corpus callosum, as demonstrated in Fig. 4h and Fig. 5h.

As FA decreases with an increasing *b*-value, as shown in Fig. 6a, and the threshold of FA for determining WM image voxels was fixed in our work, the number of selected WM image voxels will decrease with a higher *b*-value compared to that with a lower *b*-value, as demonstrated in Fig. 3. More importantly, as shown in Fig. 6d, the derived 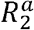 will slightly increase with a higher *b*-value because the selected WM image voxels generally possess more anisotropic biophysical features [17]. Therefore, care should be taken when comparing prior reports and the findings from the present work.

Our work relies on a single T2W image, rather than conventional *T*_2_ mapping that acquires multiple T2W images with varying echo times. Therefore, the uniformity of the T2W image plays a crucial role in deriving accurate *R*_2_ orientation dependence in WM. Any systematic errors leading to bias in the signal intensities of the T2W image will directly translate into an unreliable anisotropic *R*_2_ value. The high-resolution diffusion datasets used in the present work indeed manifested a visible asymmetry from left to right brain hemisphere on an axial imaging plane (data not shown). It remains unclear to what extent this signal asymmetry would compromise our results; however, the overall anisotropic transverse relaxation trends observed from CC1 to CC7 are expected to hold, as they were exposed to the same systemic errors. The diffusion metrics were, however, not affected by this systematic error due to the normalization process involved in DTI postprocessing.

### 5.3 Comparing with previous callosal diffusion and relaxometry

As shown in Fig. 6a, the measured FA across the corpus callosum exhibits an increasing pattern from the anterior (CC1) to posterior (CC7) segments, consistent with previous findings, particularly for the second half of the segmented tracts [44,43,46,51]. Unlike the present work, previous studies did not apply both FA and MO thresholds for callosal image voxels, which likely contributed to the observed differences between the current and prior findings. Nevertheless, an axon fiber with a larger radius seems to be associated with a higher FA value, such as in the splenium (CC7).

Recall that C_0_, as shown in Fig. 6c, was defined as 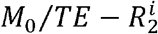. Although the relative variation is small, about 5%, the observed increasing trend of *C*_0_ would translate into an increasing trend of *M*_0_ with a constant 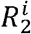 or a decreasing trend of 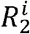 with a constant *M*_0_. We could not exclude another scenario with an increasing 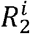, where the observed variation of *C*_0_ should be attributed mostly to *M*_0_ variations. In the literature [46], *M*_0_ was shown to increase from the anterior to posterior, equivalent to from CC1 to CC7. Derived from a mono-exponential model, 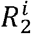 was found to decrease from CC1 to CC4 and then increase from CC5 to CC7, in a recent study on compartmental *T*_2_-orientational dependence [23].

In the past [51,46,52], the longitudinal relaxation rate *R*_1_ has been intensively investigated across the callosal segments, showing a decreasing trend from CC1 to CC7. This relaxation metric was found to be positively proportional to fiber density [52] and negatively proportional to axon size [53]. The posterior callosal segments tend to have larger-diameter axon fibers than the anterior locations according to previous light microscopic findings [54]. If we interpret this *R*_1_ decrease in terms of more restricted water molecular reorientations associated with larger-diameter axon fibers on the timescales of nanoseconds, 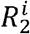 would increase across the callosal segments from CC1 to CC7.

### 5.4 Orientation-dependent *R*_2_ in WM linked to (de)myelination

When CC3 was excluded due to suboptimal fits (see Table 1), 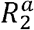 showed an increasing trend from CC1 to CC7 as illustrated in Fig. 6d. This specific relaxation metric suggests that more rotationally restricted water molecules or their host macromolecules (e.g., myelin sheath) were found in the posterior segments than in the anterior segments. Therefore, we can conclude that an increased myelination accompanies an increased axon size from CC1 to CC7. This conclusion aligns well with previous electron microscopic findings, which indicate that not only axon diameter but also myelin thickness shows an increasing trend from the anterior to the posterior callosal segments [55]. It is worth mentioning that both the derived 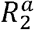 from this work and the histologically determined axon diameter and myelin thickness from the literature reached their maximum before the most posterior segment.

As shown in Fig. 6e, no obvious trend was identified for the derived angle *α*. According to our previously developed orientation-dependent *R*_2_ model [17], a constant ratio of anisotropic *R*_2_ contributions from IAS and EAS across the callosal segments will lead to a constant *α*. This biophysical interpretation of the measured anisotropic relaxation metrics is schematically depicted in Fig. 7a. In this schematic, we assumed the observed fluctuated angle *α* to be constant after excluding CC3 and considering the higher uncertainties for some *α* values.

**Fig. 7.**
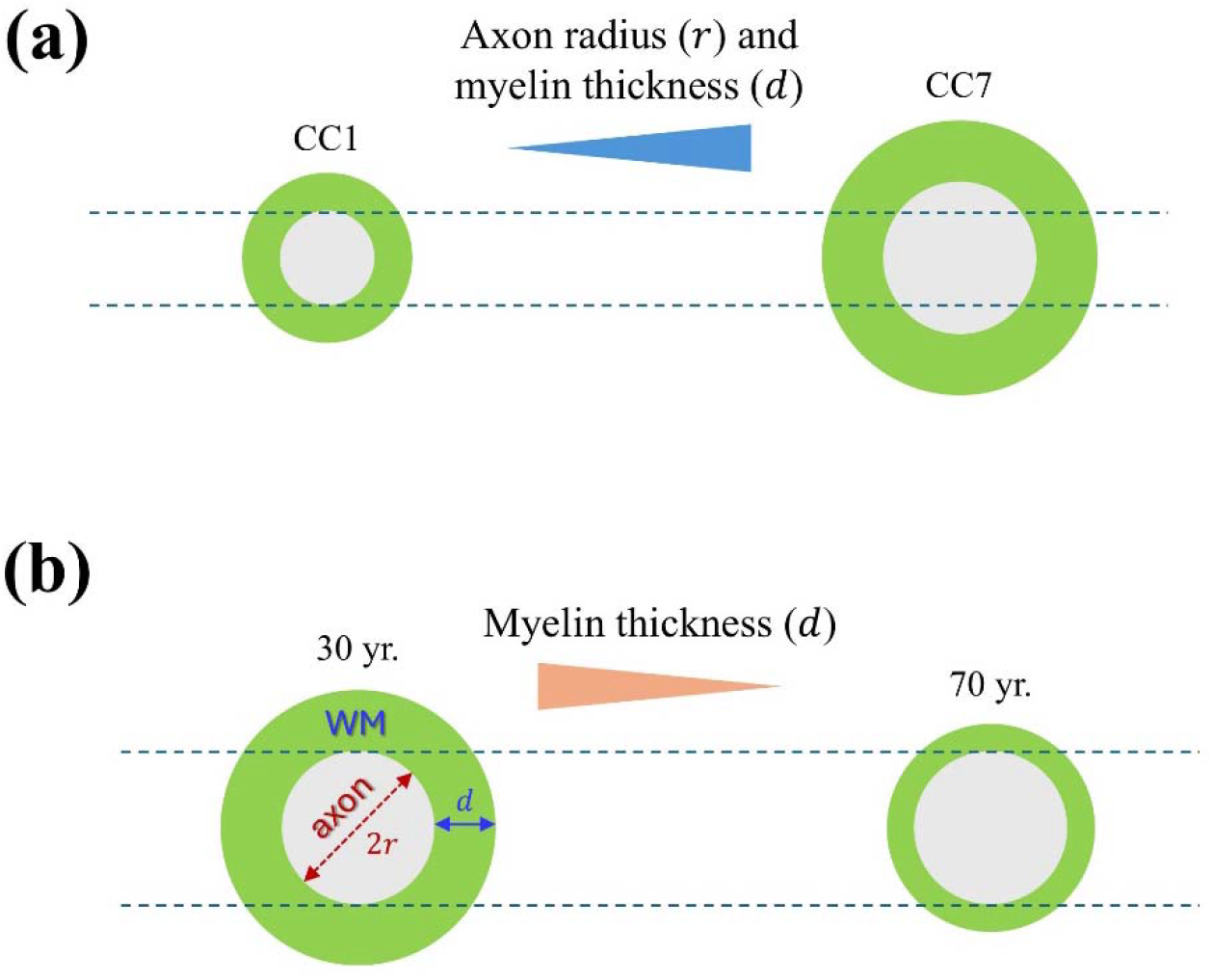
Schematic of (de)myelination with a varying axonal size (a) or a fixed axonal size (b). In this schematic, ‘r’ denotes the radius of an exemplary axon, and ‘d’ represents the thickness of the myelin sheath.

In our previous study, we showed that both 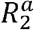 and *α*, derived from the whole brain WM, decreased with aging (Fig. 8D-E in the reference [28]). If an anisotropic *R*_2_ contribution from IAS remains constant regardless of aging, a decreasing *α*would be expected. This is a good example of clinically relevant demyelination, and its biophysical interpretation is demonstrated schematically in Fig. 7b. Consistently, the measured 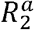 from the subjects with multiple sclerosis (MS) was much smaller compared to those from the two control groups [28,31]. If we consider the changes in 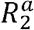 to be positively correlated with myelin content, the increase from CC1 to CC7 shown in Fig. 6d and the decrease with aging, as shown previously, are consistent.

Because of the interplay between 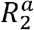 and *α* in terms of WM microstructural alterations, cautions should be exercised when interpreting the measured 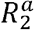. Based on the developed cone model [28,17], 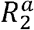 comprises two contributions: one from the extent of myelination, assuming that the residual dipolar vectors of the rotationally restricted water molecules are perpendicular to the axon fiber direction (i.e., *α*= 90°), and the other from within the axonal space, with the residual dipolar vectors assumed to be aligned with the axon fiber direction (i.e., *α*=0°) [25].

Therefore, the same increase in 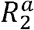 might be induced by different microstructural changes, depending on the measured open angle *α* of the cone model, as schematically shown herein in Fig 7. More specifically, while both scenarios in Fig. 7a from CC7 to CC1 and Fig. 7b from the young to the old could be considered as demyelination cases, the former could also be associated with axonal loss concurrent with demyelination.

### 5.5 Limitations

Although the proposed method is straightforward and can be easily applied to clinical DTI applications, we must point out its limitations in practical uses. First, the derived cone model parameters depend critically on the criteria for selecting WM voxels. It has been shown that the extent of the orientation dependent *R*_2_ can be dramatically reduced when the threshold of fractional anisotropy, a selection criterion, is lowered [56]. Moreover, fractional anisotropy depends on the *b*-value for diffusion weightings, as shown in Fig. 6a. Second, the quality of T2W images could be negatively impacted by any potential inhomogeneous *B*_1_ fields, which could directly translate into biased model parameters derived from non-uniform T2W images. Third, in this work, the analysis focused only on the segmented fiber tracts passing through the corpus callosum. These segmented callosal tracts provide sufficient orientation samples that span the full angular range, making data modeling feasible. For other fiber bundles with limited orientation samples, the presented method will not work, and an alternative approach should be sought if specific WM microstructural information needs to be disentangled. Such an alternative could be *R*_1*ρ*_ dispersion, as previously demonstrated in articular cartilage [32]. Finally, no effort has been made to estimate the extent to which the “susceptibility” effect may have influenced the derived 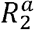 in this work. However, previous research [57–59] and our latest findings (to be published) inferred from 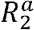 measurements at 3 T and 7 T lead to the conclusion that the measured 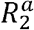 at 3 T is predominately induced by the magic angle effect, consistent with the literature [28].

## 6. CONCLUSIONS

In conclusion, we have demonstrated that fiber-tract-specific anisotropic *R*_2_ profiles can be conveniently derived from DTI and further characterized using an established cone model. The proposed method offers a unique opportunity to reevaluate existing clinical DTI studies and optimize new ones for characterizing (de)myelination in both healthy and diseased subjects.

## ACKNOWLEDGEMENT

We would like to thank Dr. Fuyixue Wang (Massachusetts General Hospital, Charlestown, MA, USA) for sharing Connectome high-resolution DTI datasets in the public domain. The authors also thank the anonymous reviewers for their insightful comments and helpful suggestions. This work has been enhanced with the assistance of Microsoft Copilot for proofreading and quality improvement.

## AUTHORS’ CONTRIBUTIONS

Pang: contributed to study conception and design, analysis and interpretation, drafting of manuscript, and critical revision. Raja: contributed to data analysis and critical revision. Reddick: contributed to data interpretation and critical revision.

## DATA AVAILABILITY STATEMENT

Data and analysis software (Python and IDL) used in this work are available on request from the authors.

## DECLARATION OF COMPETING INTERESTS

The authors have no conflicts of interest to declare.

## FUNDING INFORMATION

This work was funded, in part, by the American Lebanese Syrian Associated Charities (ALSAC) at St. Jude Children’s Research Hospital.

## Notes

### Competing Interest Statement

The authors have declared no competing interest.

